# An endometrial organoid model of *Chlamydia*-epithelial and immune cell interactions

**DOI:** 10.1101/2020.07.29.226969

**Authors:** Lee Dolat, Raphael H. Valdivia

## Abstract

Our understanding of how the obligate intracellular bacterium *Chlamydia trachomatis* reprograms the cell biology of host cells in the upper genital tract is largely based on observations made in cell culture with transformed epithelial cell lines. Here we describe a primary spherical organoid system derived from endometrial tissue to recapitulate epithelial cell diversity, polarity, and ensuing responses to *Chlamydia* infection. Using high-resolution and time-lapse microscopy, we catalogue the infection process in organoids from invasion to egress, including the reorganization of the cytoskeleton and positioning of intracellular organelles. We show this model is amenable to screening *C. trachomatis* mutants for defects in the fusion of pathogenic vacuoles, the recruitment of intracellular organelles, and inhibition of cell death. Moreover, we reconstructed a primary immune cell response by co-culturing infected organoids with neutrophils, and determined that the effector TepP limits the recruitment of neutrophils to infected organoids. Collectively, our model details a system to study the cell biology of *Chlamydia* infections in three dimensional structures that better reflect the diversity of cell types and polarity encountered by *Chlamydia* upon infection of their animal hosts.

**Summary statement:** 3D endometrial organoids to model *Chlamydia* infection and the role of secreted virulence factors in reprogramming host epithelial cells and immune cell recruitment

## INTRODUCTION

*Chlamydia trachomatis* is a clinically important pathogen, responsible for the majority of sexually transmitted bacterial infections (WHO, 2018). In the female upper genital tract (UGT), chronic and untreated asymptomatic infections can lead to pelvic inflammatory disease, tubal scarring and infertility (Haggerty et al., 2010), and are associated with cancers of the cervix and endometrium (Koskela et al., 2000). Yet, despite decades of studies the molecular mechanisms by which *C. trachomatis* disrupts epithelial tissue structure and induces pathology in the UGT are not well understood.

The genus *Chlamydia* is comprised of nine *Chlamydia* species that infect eleven different vertebrate hosts, including the human pathogen *C. trachomatis* and rodent-adapted *C. muridarum* (Horn, 2008; Schachter, 1978). Based on sequence similarity and disease manifestations, *C. trachomatis* is further classified into serovars and biovars that target the genital tract (serovars D-K) and ocular epithelia (serovars A-C), the etiological cause of the inflammatory disease trachoma (Stephens et al., 2009). The lymphogranuloma venereum (LGV) biovar (serovars L1-L3) is an invasive infection in the urogenital or anorectal tracts that disseminates to the lymph nodes (Mabey and Peeling, 2002). Of all the serovars, *Chlamydia* LGV serovar L2 is the only one that is fully tractable to molecular genetic manipulations (Mueller et al., 2016; Wang et al., 2011).

All *Chlamydia* transition between two developmental forms: the environmentally stable and infectious elementary body (EB), and the non-infectious but replication-competent reticulate body (RBs) (Abdelrahman and Belland, 2005). To invade cells, EBs use a type III secretion system (T3S) to deliver effector proteins directly into host epithelial cells. These invasion-associated early T3S effectors stimulate rearrangements in filamentous actin (F-actin) at attachment sites to promote entry and uptake into an intracellular membrane vacuole termed the “inclusion.” (Elwell et al., 2016). Additional effectors are then secreted and inserted into the nascent inclusion membrane. These effectors, termed inclusion membrane (Inc) proteins, co-opt microtubule-based trafficking to aid in the migration of inclusion to the microtubule organizing center, subvert intracellular organelles, including the Golgi apparatus, endoplasmic reticulum, peroxisomes, and lipid droplets, and establish membrane contact sites to presumably facilitate the acquisition of nutrients form its host (Agaisse and Derré, 2014; Boncompain et al., 2014; Derré et al., 2011; Dumoux et al., 2012; Grieshaber et al., 2003; Kumar et al., 2006; Mital et al., 2015; Moore et al., 2008). Within the inclusion, EBs differentiate into RB forms, replicate and expand along the perimeter of the inclusion. The inclusion itself is wrapped in a network of actin and intermediate filaments, microtubules, and septins (Dumoux et al., 2015; Kumar and Valdivia, 2008; Volceanov et al., 2014; Wesolowski et al., 2017). RBs asynchronously differentiate back into EBs and escape from the cell through a F-actin-mediated extrusion of the inclusion or host cell lysis to initiate the next round of infections (Hybiske and Stephens, 2007).

*Chlamydia* maintains an intracellular replicative niche by suppressing cell-autonomous innate immunity (Finethy and Coers, 2016). For example, *Chlamydia*-driven rearrangements of the cytoskeleton promotes inclusion stability, limiting its detection and activation of the innate immune response (Kumar and Valdivia, 2008). *Chlamydia* Inc proteins counteract host defenses by deubiquitinating proteins at the inclusion, inhibiting the lytic activity of interferon induced factors (e.g. guanylate binding proteins) and suppressing cell death by subverting membrane trafficking to inhibit host surveillance programs and apoptosis (Coers et al., 2008; Faris et al., 2019; Fischer et al., 2017; Sixt et al., 2017). Despite these activities, *Chlamydia* is still recognized by the host cell which can lead to a pro-inflammatory response (Rasmussen et al., 1997), including the expression and secretion of chemokines, such as the neutrophil attractants interleukin-6 (IL-6) and interleukin-8 (IL-8) (Buchholz and Stephens, 2008; Rasmussen et al., 1997).

Our understanding of the cell biology of how *Chlamydia* re-programs host cellular functions is based largely on studies in two-dimensional cell culture systems with transformed cervical epithelial cell lines like HeLa cells. While these models are experimentally tractable and have been of great use, they do not accurately represent the natural sites of infection: polarized UGT epithelial cells that have a distinct spatial organization of the cytoskeleton, organelle positioning, and signaling modules that are distinct from transformed nonpolarized epithelia (Dolat and Valdivia, 2019; Rodriguez-Boulan and Macara, 2014). Infections in polarized epithelia have uncovered unique aspects of the interaction between *Chlamydia* and its target cell: *C. trachomatis* replication is enhanced in polarized epithelial cells and the inclusion preferentially intercepts vesicles destined for the basolateral side of infected cells (Guseva et al., 2007; Moore et al., 2008). Infections in primary polarized human ecto- and endocervical epithelial explants revealed that *Chlamydia* alters epithelial structure by inducing epithelial-to-mesenchymal transition (Zadora et al., 2019). Furthermore, long-term *Chlamydia* infections in fallopian tube organoids, self-renewing primary three-dimensional epithelial tissue-like cultures, promotes epithelial stemness, cellular proliferation, and modifications to genome organization often associated with ageing tissues (Kessler et al., 2019). These observations may help explain some of the associations observed between *Chlamydia* infections and cervical and ovarian cancers (Shanmughapriya et al., 2012; Zhu et al., 2016).

Three-dimensional organoids have emerged as a compelling *ex-vivo* system to study epithelial cell biology as they better recapitulate aspects of the cellular diversity, structure and function of epithelial tissues (Rossi et al., 2018). Isolated human and mouse glandular endometrial epithelia cultured in a chemically defined media were recently reported to form endometrial organoids (EMOs) comprised of hormonally responsive polarized epithelial subtypes (e.g. ciliated, secretory) that mimic aspects of the estrous cycle and implantation (Boretto et al., 2017; Turco et al., 2017). EMOs exhibit long-term genetic stability and are amenable to high-throughput screens (Boretto et al., 2019). Here, we describe an infection model that uses EMOs and microinjection to deliver *Chlamydia* to the apical surface of polarized epithelium, thus mimicking the natural route of infection. Using both the rodent-adapted *C. muridarum* and the genetically-tractable human pathogen *C. trachomatis* serovar L2, we demonstrate key aspects of the infection cycle and cellular activities driven by *C. trachomatis* effectors. We further show that we can recapitulate aspects of cellular immunity in infected EMOs by monitoring the recruitment and behavior of neutrophils. Using a *C. trachomatis* T3S effector mutant that alters immunity-related signaling pathways, including the expression of neutrophil chemoattractants, we further demonstrate that this system can be used to measure the manipulation of host immune responses by *Chlamydia* virulence factors.

## RESULTS

### Cells harvested from the upper genital tract develop into endometrial organoids of cellular composition similar to that of primary tissues

Isolated endometrial glandular epithelia were cultured in a three-dimensional Matrigel matrix in the presence of conditioned medium from L-WRN cells. L-WRN cells stably express and secrete Wnt-3A, R-Spondin 3, and Noggin, efficiently generate organoids from various mouse tissues with little batch-to-batch variation (Miyoshi and Stappenbeck, 2013; VanDussen et al., 2019). Indeed, EMOs readily form over the course of one week and grow into large, spherical structures (Fig 1 B-C).

**Figure 1.**
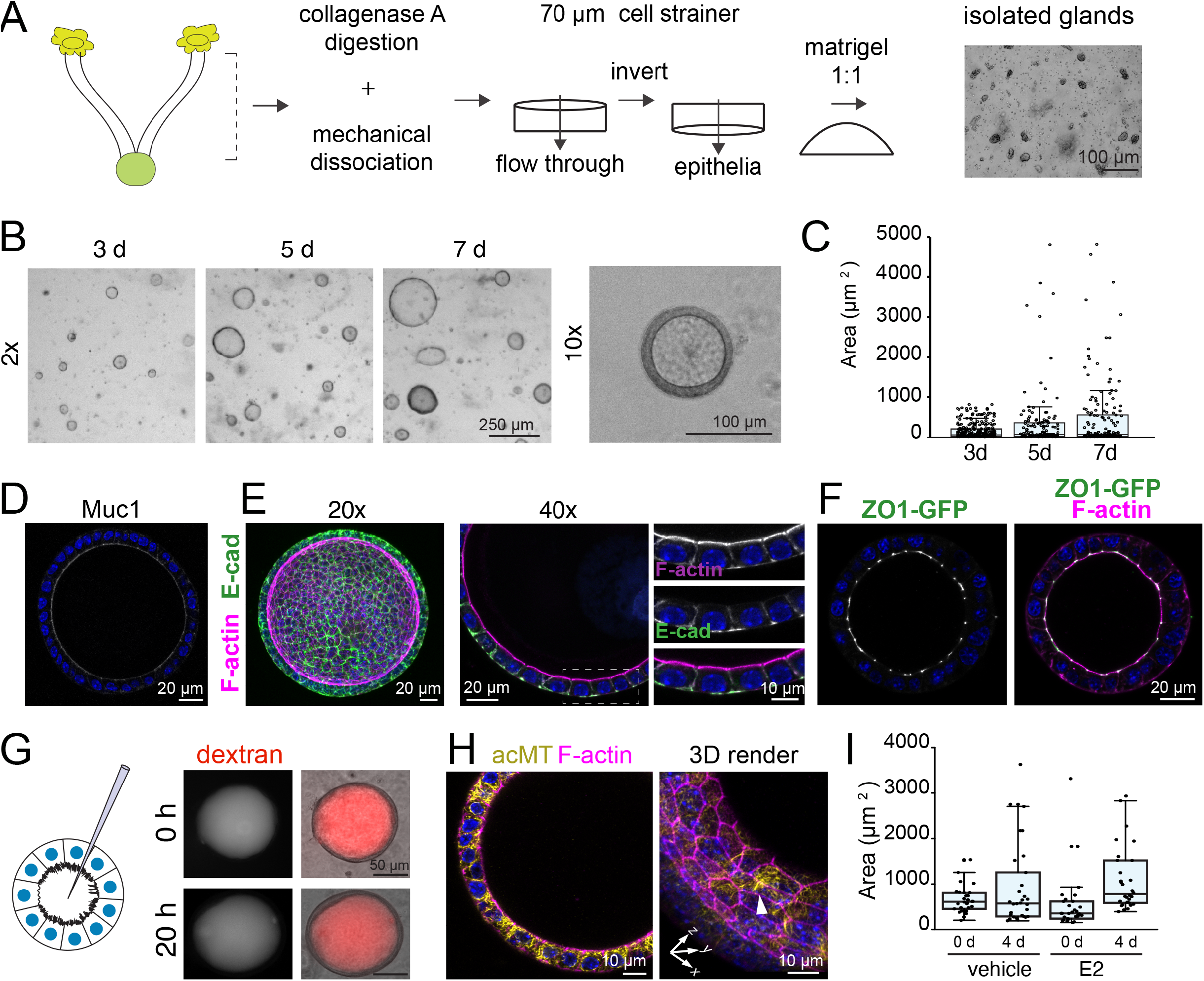
Generation of primary murine-derived polarized endometrial epithelial organoids. (**A**) Workflow for the isolation of endometrial epithelial glands and culture in three-dimensional extracellular Matrigel plug. (**B-C**) Cells in endometrial organoids (EMOs) proliferate in the presence of Wnt-3a, R-Spondin3, Noggin, and EGF. Brightfield images of organoids forming over a seven day period are shown, including a single EMO at higher magnification and quantification of the relative surface area (n > 100 EMO per timepoint) (C). (**D-F**) EMO express Muc1 and establish tight and adherens junctions. (D) Immunofluorescence microscopy of Muc1 (white) in EMOs. (E) Immunofluorescence microscopy of the epithelial marker E-cadherin. Left: Polarized EMOs established adherens junctions. Confocal images of EMOs stained for DNA (blue), F-actin and E-cadherin using a 20x objective (left, maximum projection) and a 40x objective (right, medial section). (F) EMOs establish tight junctions. Medial section of confocal microscopy images of a ZO1-GFP knock-in EMO stained for F-actin and DNA (blue). **(G)** EMO epithelia maintain barrier function. *Left;* Cartoon of endometrial organoid microinjection system. *Right;* EMO microinjected with Texas-Red dextran and imaged immediately post-injection and 20 hours later. **(H)** Immunofluorescence microscopy of an EMO treated with estrogen for four days. Confocal images an EMO stained for acetylated-tubulin, F-actin, and DNA (blue). Arrow points to multi-ciliated epithelial cell. **(I)** Estrogen promotes EMO growth. Quantification of EMO size with and without estrogen (E2) for four days (n = 30 EMOs per condition).

Using high-resolution confocal microscopy, we determined that EMOs are comprised of a single layer of polarized epithelial cells surrounding a large, hollow lumen. The epithelia express and secrete mucin (Muc1) (Fig 1D) and establish adherens and tight junctions (Fig 1E-F), respectively marked by E-cadherin and zonula occludens-1 (ZO-1). To assess the integrity of the spheroids and the functionality of the cell-cell junctions, we microinjected fluorescent 3 kDa dextran into the organoid lumen. The fluorescent tracers remained in the lumen indicating that barrier function is maintained over prolonged periods of time (Fig 1G). Endometrial tissues are responsive to sex hormones, increasing cellular proliferation and altering gene expression and cell type abundance in the presence of estrogen and progesterone. EMOs are similarly responsive as assessed by the larger increase in size upon exposure to estrogen for four days, as compared to untreated EMOs (2.0x vs 1.37x), and the formation of multi-ciliated epithelia as marked by acetylated microtubules (Fig 1H-I). Overall, the method produces hormonally responsive polarized EMOs with intact barrier function and a distinct lumen.

### *Chlamydia* infection of endometrial epithelial leads to cytoskeletal reorganization and disruption of cell-cell junctions

To mimic the natural route of *Chlamydia* invasion we microinjected GFP-expressing *C. trachomatis* serovar L2 or *C. muridarum* CM006, a high pathology isolate (Poston et al., 2018), into the EMO lumen. We monitored the subcellular localization of F-actin and β-catenin, which are known to localize to the inclusion (Kessler et al., 2012) at early timepoints (8-16 hours) post injection. EBs readily invaded the polarized epithelial cells, promoted the assembly of actin filaments as the nascent inclusion formed, and recruited β-catenin (Fig 2A-B). Consistent with a previous ex vivo fallopian tube infection models (Kessler et al., 2012), infected epithelia showed a striking loss of apical F-actin and cell-cell junction integrity, demarcated by more diffusive β-catenin localization, indicating that epithelial polarity and barrier function are compromised during invasion. We observed β -catenin recruitment to inclusions only at very early time points with no significant recruitment to inclusions by 48 hpi (Fig S1A-B). More recently, *Chlamydia* has been shown to target epithelial tight junctions, altering the expression of tight junction proteins and reducing transepithelial electrical resistance (Kumar et al., 2019). Using EMOs derived from ZO1-GFP expressing transgenic mice (Foote et al., 2013), we show that ZO-1 is recruited to EBs during invasion along with β-catenin, which may further underlie the disruption to epithelia barrier functions (Fig 2C). Taken together, these data show that *Chlamydia* invasion remodels the F-actin network and cell-cell junction organization in a manner that disrupts cell polarity early during the infectious process.

**Figure 2.**
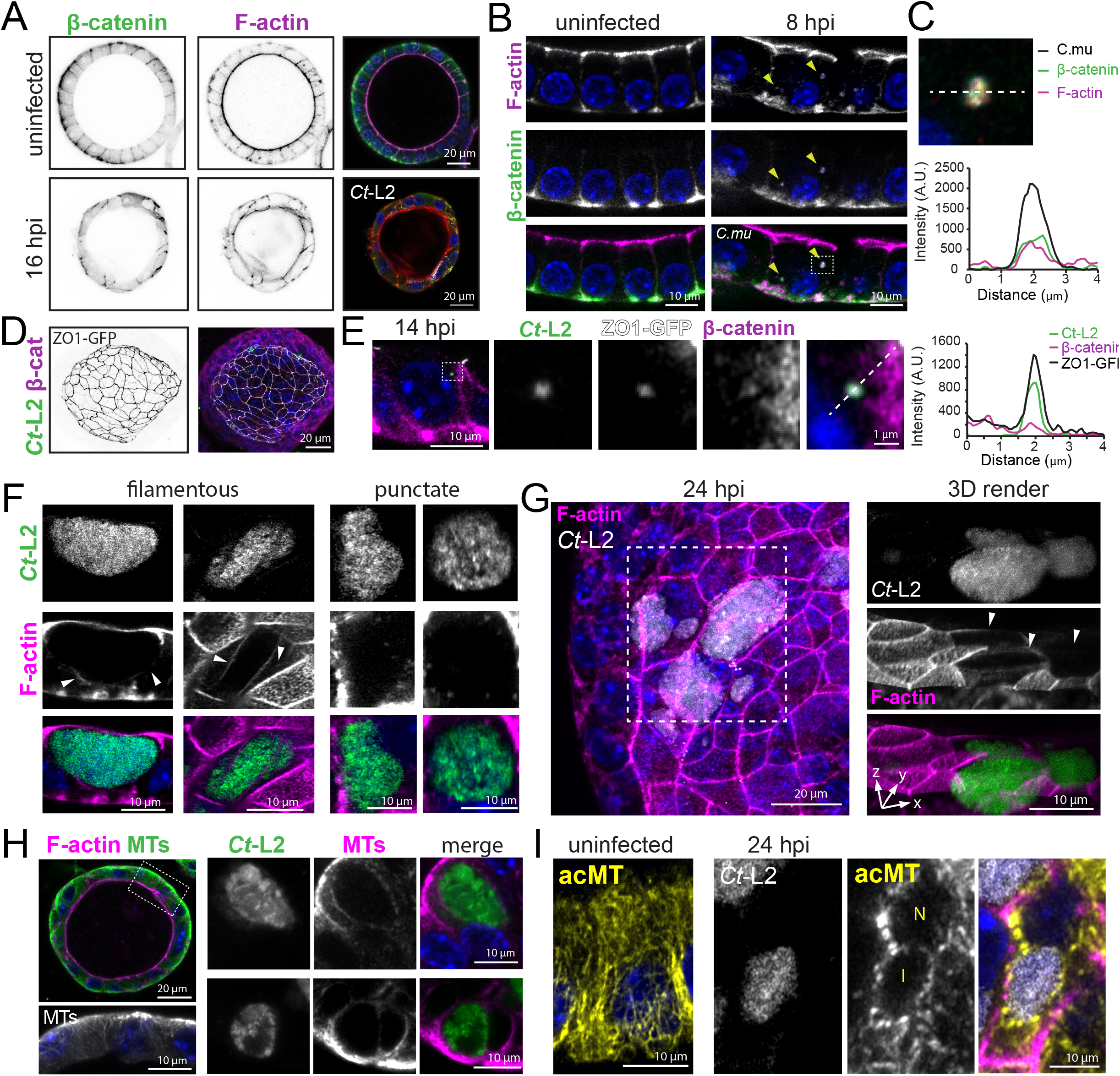
*Chlamydia* infection leads to the remodeling of the endometrial epithelial cytoskeleton and cell-cell junctions. (**A-C**) *C. trachomatis* (**A**) and *C. muridarum* (**B**) infection disrupt endometrial epithelial cell-cell junction organization. Confocal images of EMOs uninfected or infected with for 14 h (*Ct*-L2) or 8 h (*C. mu*) and stained for F-actin, β-catenin, and DNA (blue). GFP-expressing *C. muridarum* are marked by yellow arrowheads with a fluorescence line scan profile of a single inclusion showing the co-recruitment of F-actin and β-catenin (**C**). (**DE**) *C. trachomatis* early inclusions recruit the tight junction protein ZO-1. Maximum projection confocal images of EMOs derived from ZO1-GFP expressing mice before (**D**) and single confocal slices after infection with mCherry-expressing *Ct*-L2 for 14 h (**E**) and stained for β-catenin and DNA (blue). Enlarged panels display a single inclusion and the recruitment of ZO-1 and β-catenin and fluorescence line profile of *Ct*-mCherry, ZO-1, and β -catenin. (**F-G**) *C. trachomatis* reorganizes the cytoskeleton of EMO cells. Confocal images of EMOs infected with GFP-expressing *Ct*-L2 for 24 h and stained for F-actin (**F**). *Left panels*: filamentous actin ring-like structures and/or cages around the inclusion. *Right panels*: F-actin puncta around the inclusion. *C. trachomatis* reduces F-actin signal on the apical membrane (**G**) as assessed by 3D rendering (boxed region) of *Ct*-L2 (GFP) infected cells (24 h) in EMOs after staining for F-actin. Arrowheads point to infected cells lacking apical F-actin. (**H-I**) Organization of microtubules (MTs) in EMOs. Confocal images of an uninfected EMO stained for microtubules (MT) and F-actin (H) and enlarged images of microtubules rearranged at the periphery of *Ct*-L2 inclusions at 24 hours post-infection. *C. trachomatis* promotes the assembly of stable microtubules around the inclusion in EMO as assessed by immunostaining for acetylated microtubules (I). Confocal images of acetylated microtubules in an uninfected epithelial cell (*left panel*), and an EMO infected with GFP-expressing *Ct*-L2 for 24 hours (*right panels*). For all images DNA stains used 2 μg/mL Hoechst (blue).

During the mid and late stages of infection *Chlamydia* remodels the cytoskeleton of HeLa cells at the periphery of inclusions (Kumar and Valdivia, 2008). We monitored the localization of various cytoskeletal elements in infected EMOs at mid-cycle (24 hpi) by probing for F-actin and immunostaining against microtubules, cytokeratins, and septins – all of which localize to the inclusion periphery (Chin et al., 2012; Kumar and Valdivia, 2008; Volceanov et al., 2014). In polarized epithelial the cytoskeletal architecture is markedly different from non-polarized and transformed cells. For example, the F-actin network is enriched at the apical membrane, microvilli, and cell-cell junctions rather than a dense network of bundled actin stress fibers (Fig 2A). Moreover, in contrast to 2D culture settings, microtubule organizing centers localize proximal to the apical membrane and generate microtubules that extend towards the basolateral domain, often along the lateral membrane (Pickett et al., 2019). We observed F-actin structures around the inclusion, either as filamentous-like rings or discrete puncta, which may depend on the stage of the infection cycle (Fig 2E). Moreover, F-actin signal on the apical membrane was markedly reduced or absent in infected epithelia compared to uninfected neighboring cells (Fig 2F), suggesting that infection and/or inclusion growth promotes the loss of microvilli. In uninfected EMOs, microtubules run along the apico-basolateral axis and terminate at the basolateral membrane (Fig 2G). During infection, we observed microtubules and acetylated microtubules prominently assembled around the inclusion (Fig 2H-I). Because *Chlamydia* effectors, such as CT288 and IPAM, target centrosomal proteins (Almeida et al., 2018; Dumoux et al., 2015), this model can be used to explore effector functions in MT assembly during infection.

The organization and dynamics of F-actin and microtubules are regulated directly by septins, a conserved family of GTPases that assemble into heteropolymers and filaments (Mostowy and Cossart, 2012; Spiliotis, 2018). In EMOs, septin 2 (Sept2) localizes to the apical and lateral membranes and partially co-localizes with F-actin at the basolateral surface (Fig S1C). In infected EMOs Sept2 undergoes a dramatic reorganization to forms filamentous rings around the inclusion (Fig S1D) as had been previously observed in non-polarized cells (Volceanov et al., 2014).

Finally, we probed the endometrial epithelia for intermediate filaments. Both keratins and vimentin localize around the inclusion in transformed epithelia and their absence disrupts inclusion stability (Kumar and Valdivia, 2008; Tarbet et al., 2018). As expected, EMO epithelia express keratins (Fig S1E) which relocalize to the *C. trachomatis* inclusion (Fig S1F). However, unlike transformed epithelia, EMO epithelial cells do not express vimentin (Fig S1G), and its expression is not induced during infection with *C. muridarum* (Fig S1H). In contrast, primary endometrial stromal fibroblasts infected with *C. muridarum* and *C. trachomatis* displayed prominent vimentin filaments cages surrounding inclusions (Fig S1I).

Overall these findings indicate that despite the difference in cytoskeletal organization between transformed, non-polarized epithelial cells, *Chlamydia* likely employs conserved mechanism to reorganize these structures during cell entry and formation of inclusions.

### Live imaging of inclusion expansion, dynamics and exit

We developed an imaging platform to monitor inclusion growth, dynamics and exit using timelapse 3D spinning disk confocal microscopy. EMOs were infected for 24 hours with *C. trachomatis* serovar L2 expressing GFP and subsequently imaged live for 16-18 hours. We noted inclusion expansion and dynamics, including the apparent fusion of adjacent inclusions (Fig 3A). We next tested if inclusion fusion is a phenomena that occurs in EMOs by co-infecting with *C. trachomatis* L2 expressing GFP or mCherry, or with a *C. trachomatis* mutant (M923) lacking the fusogenic factor IncA (Kokes et al., 2015; Sixt et al., 2017). GFP and mCherry positive inclusions were readily apparent EMOs co-infected with wild-type strains (Fig 3B) but not in EMOs coinfected with the IncA mutants (Fig 3B), indicating that this model can be employed to investigate the propensity of inclusions to fuse and if this process is influenced by the cell type being infected.

**Figure 3.**
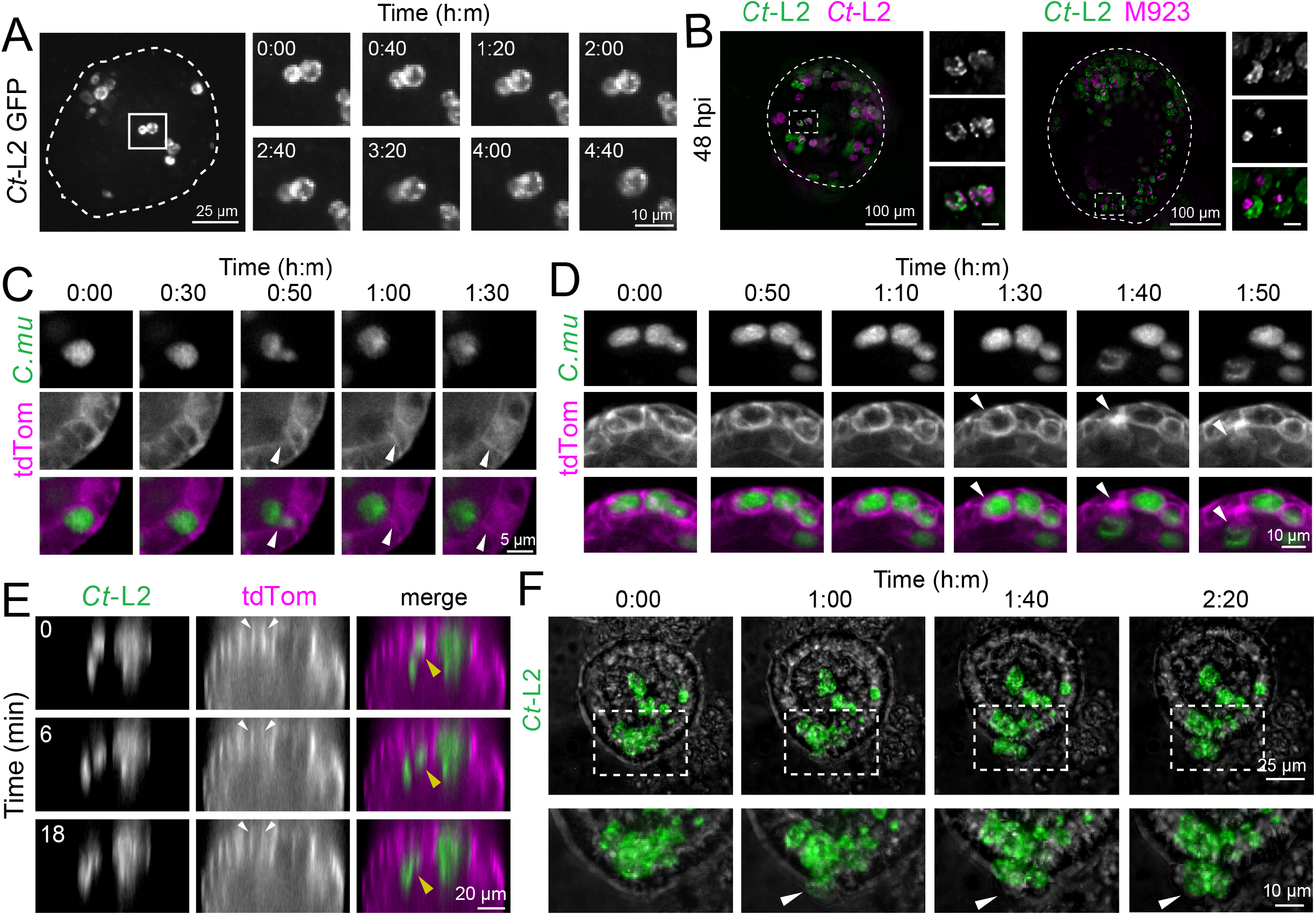
Time-lapse microscopy of inclusion dynamics and exit from infected endometrial epithelia. (**A-B**) *C. trachomatis* inclusions are fusogenic in EMO. Maximum projections of livecell fluorescence spinning disk microscopy of an EMO infected with GFP-expressing *C. trachomatis*. Enlarged frames show two inclusion fusing over time (A) and that fusion is blocked when co-infections are performed with an IncA mutant (M923) expressing mCherry. Scale bars, 10 μm. (B). EMOs were imaged by 3D widefield deconvolution microscopy. (**C-D)** Apical extrusion of *C. muridarum* inclusions and infected host cell in EMOs. TdTomato EMOs were infected with GFP-expressing *C. muridarum* for 24 hours and imaged by time-lapse 3D spinning disk confocal microscopy. Single confocal slices at the indicated timepoints show an inclusion released into the lumen (C). White arrowheads denote infected epithelial cell that remains intact, and lysis of the infected epithelial cell along with the inclusion into the lumen (D). (**E-F**) *C. trachomatis* L2 inclusions extrude apically and basolaterally. (E) Orthogonal view of a tdTomato EMO infected with GFP-expressing *Ct*-L2 and imaged by time-lapse spinning disk microscopy. Yellow arrowhead denotes an inclusion extruding into the inclusion lumen and white arrowheads indicate intact cell boundaries. (F) Maximum projections from time-lapse spinning disk microscopy of an EMO infected with GFP-expressing *Ct*-L2. White arrowheads denote cells/inclusions that bulge basolaterally into the matrix.

We next monitored inclusions exit dynamics using EMOs derived from ROSA^mTmG^ mice that express a membrane-targeted tdTomato allowing for visualization of the plasma membrane of individual cells. First, EMOs were infected with either GFP-expressing *C. muridarum* for 24 hours or *C. trachomatis* L2 for 48h and imaged live for an additional 16 hours. We observed that *C. muridarum* inclusions were released exclusively into the organoid lumen via lysis or extrusion, which left the infected cell intact, suggesting that apical exit may be the natural mechanism by which *C. muridarum* escapes the host cell (Fig 3 D-E). In contrast, *C. trachomatis* L2 inclusions were observed to extrude both apically into the lumen (Fig. 3F) and basolaterally into the extracellular matrix (Fig. 3G).

### The reorganization of the Golgi apparatus in infected endometrial epithelia requires the inclusion membrane protein InaC

Host cell organelles are recruited to *Chlamydia* inclusions to form tethering complexes and to intercept lipid-rich vesicles and acquire nutrients (Elwell and Engel, 2012). The inclusion membrane proteins IncD, InaC, and Cdu1 have been determined to target Golgi-resident proteins and potentially regulate/intercept polarized trafficking (Agaisse and Derré, 2014; Derré et al., 2011; Kokes et al., 2015; Pruneda et al., 2018). During epithelial polarization, the Golgi apparatus migrates and expands above the nucleus, proximal to the apical membrane, where it regulates the polarized sorting of vesicles and proteins to distinct membrane domains (Bacallao et al., 1989). Using high-resolution confocal microscopy, we observed that the Golgi apparatus is more widely dispersed in EMO epithelial cells than in HeLa cells, often surrounding the nucleus and extending toward the apical surface (Fig. 4A). Consistent with previous studies (Heuer et al., 2009; Kokes et al., 2015; Pokrovskaya et al., 2012; Pruneda et al., 2018; Rejman Lipinski et al., 2009), however, we found that during early to mid-infection inclusions recruit and reorganize the Golgi apparatus (Fig. 4B).

**Figure 4.**
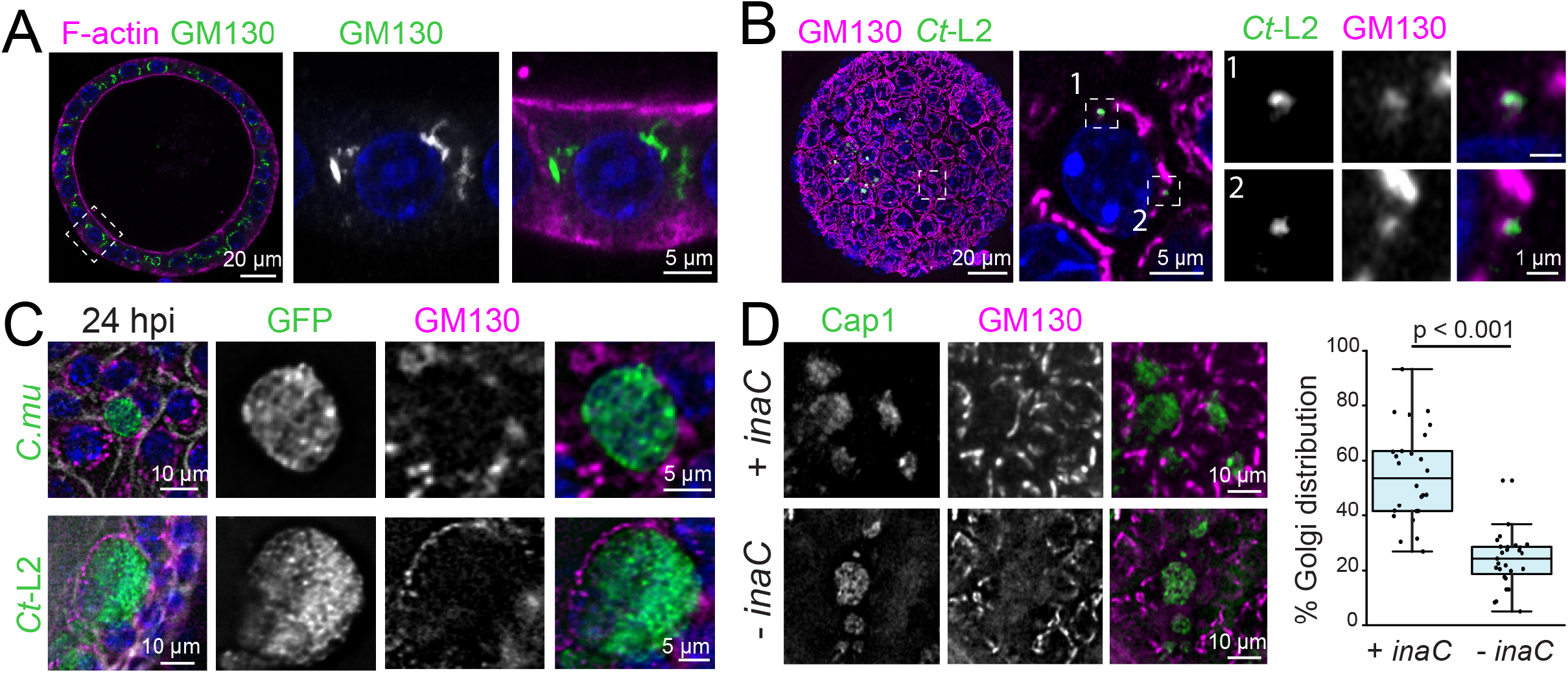
*Chlamydia* infections causes reorganization of the Golgi apparatus in endometrial epithelia. (**A-B)** The organization of the Golgi apparatus in uninfected EMOs (A) or in an EMO infected with *C. trachomatis* L2 (B). Confocal microscopy of an infected EMO stained for F-actin and GM130, as a Golgi marker. (A) Maximum intensity projection for an EMO infected with GFP-expressing *Ct*-L2 and stained for GM130. Enlarged region of interest shows relative localization of GM130 with respect to small, early inclusions in a single confocal section (B). **(C)** The Golgi apparatus relocalizes around *C. muridarum* and *C. trachomatis* inclusions at mid infection cycle. 3D deconvolution microscopy of EMOs infected with *C. muridarum* (top) or *C. trachomatis* L2 (bottom) for 24 h and stained for GM130. (**D**) InaC is required for Golgi repositioning in infected EMO cells. EMOs were infected with an *inaC* mutant M407 complemented with a plasmid expressing InaC (+ *inaC*) or an empty vector (− *inaC*) for 24 h and the localization with respect to inclusion quantified (n = 26-27 inclusions). Statistical significance was measured using a Student’s two-tailed t-test. Golgi positioning in all figures was determined by immunostaining with anit-GM130 antibodies. For all images DNA stains used 2 μg/mL Hoechst (blue).

We next visualized the extent to which the Golgi apparatus is re-positioned around the *C. muridarum* and *C. trachomatis* inclusions. Indeed, both species recruit the Golgi around the inclusion periphery to a similar extent (Fig. 4C). Because the recruitment of Golgi stacks to the inclusion is regulated by the expression and secretion of Inc proteins, such as InaC, we sought to determine if this phenotype is recapitulated in InaC mutants. EMOs were infected with the *C. trachomatis* mutant M407, bearing an InaC truncation (InaC^Q103^*), M407 complemented with *inaC* or an empty vector control (Kokes et al., 2015). Indeed, the InaC-deficient *C. trachomatis* failed to promote the redistribution of Golgi around the inclusion of infected EMO cells, while the complemented mutant shows similar distribution to that of the wild-type infected EMO cells (Fig. 4D). Collectively, these data show that despite the difference of Golgi complex morphology in polarized EMO epithelia, this organelle is still repositioned at the inclusion periphery and that this process is mediated by the same effectors.

### CpoS provides protection from *Chlamydia*-induced cell death in endometrial epithelia

Pathogen-mediated remodeling of organelle dynamics and organization can promote infection by blocking cell-autonomous immunity. For example, host recognition of microbial DNA can induce the translocation of STING (stimulator of interferon genes) from the ER to the Golgi, where it activates the interferon response (Ishikawa et al., 2009). The type I interferon response in *Chlamydia-infected* cells requires STING (Barker et al., 2013; Prantner et al., 2010), and the *Chlamydia* Inc CpoS functions to dampen the interferon response by blocking STING translocation and cell death (Sixt et al., 2017). CpoS is among a subset of Incs recently identified to promote inclusion integrity and the viability of infected cells (Sixt et al., 2017; Weber et al., 2017).

To monitor the degree of cell death in infected EMOs, we microinjected EMOs with *C. trachomatis* L2 in the presence of propidium iodide (PI) and imaged live at 24 hpi. In both uninfected and infected EMOs we observed little to no PI staining, indicating that *Chlamydia* efficiently blocks the induction of any cell death mediated defense mechanisms (Fig. 5A). We next tested if *C. trachomatis* deficient in the pro-survival factor CpoS would lead to cell death by infecting EMOs with the *C. trachomatis* mutant M007, encoding a truncated CpoS^Q31^*. Indeed, endometrial epithelial cells containing CpoS-deficient inclusions were frequently positive for PI, indicating the activation of cell death pathways (Fig. 5B). We measured the extent of this phenotype by quantifying PI signal frequency in each infected EMO and show a consistent increase with the CpoS-deficient strain (Fig. 5B-C). These observations highlight that the molecular mechanisms used by *Chlamydia* to protect infected cells from cell death-based antimicrobial responses in conserved in endometrial epithelial cells.

**Figure 5.**
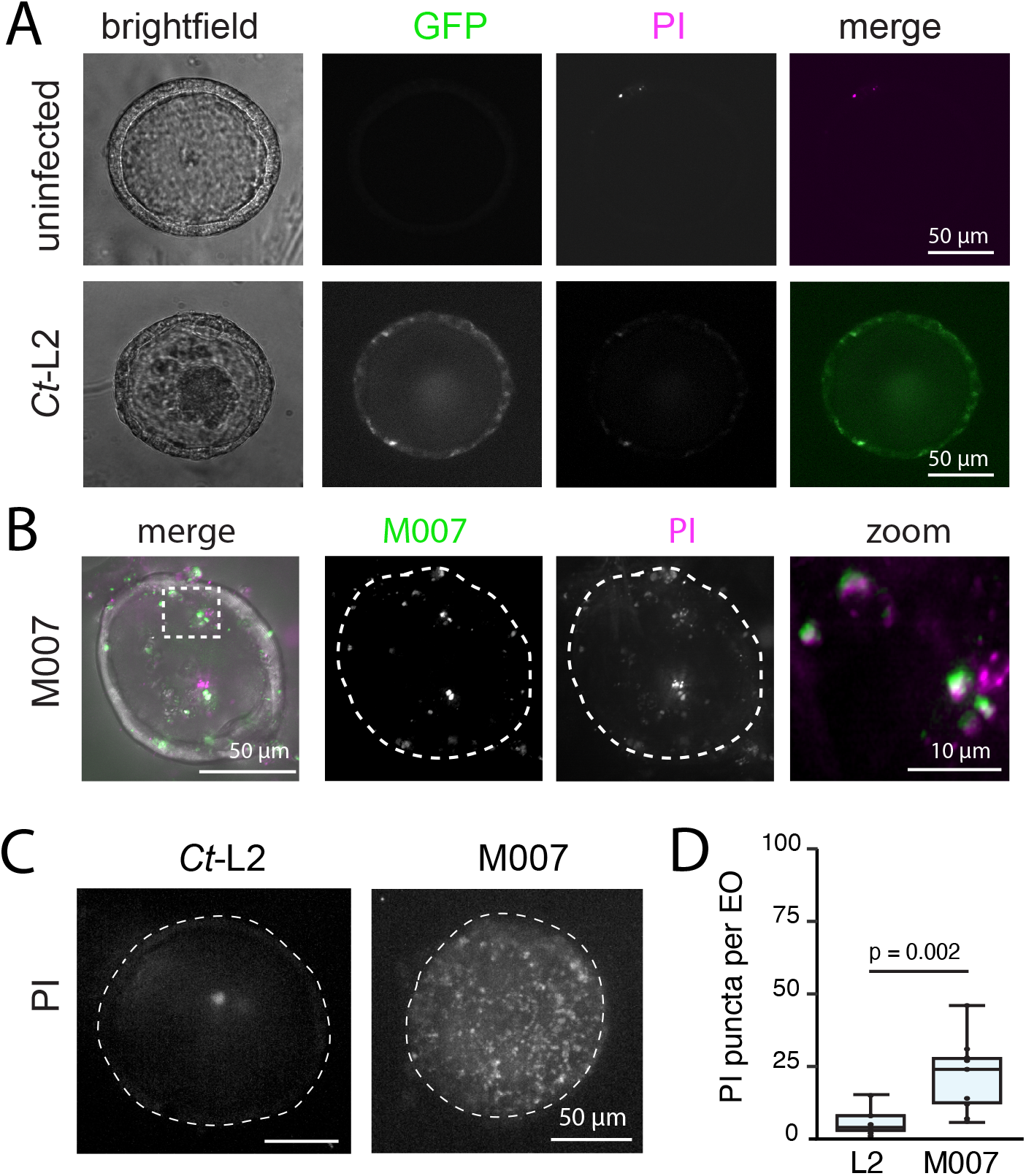
The *C. trachomatis* effector CpoS blocks cell death in infected endometrial epithelia. (**A**) EMO cells remain viable after *C. trachomatis* infection. Live spinning-disk confocal images of an uninfected EMO and an EMO infected with GFP-expressing Ct-L2 for 24 hours in media containing propidium iodide (PI). (**B-D**) EMOs infected with a *C. trachomatis cpoS* induce cell death. EMOs were infected with a GFP-expressing *cpoS* mutant (M007), incubated with propidium iodide for 22 hours, and imaged by live 3D deconvolution microscopy (B-C). Quantitative analysis of cell death, as assessed by PI-positive staining, in EMOs infected with wild-type Ct-L2 or *cpoS* M007. (C) Maximum projection images from 3D spinning-disk confocal images. Scatter plots (D) represent the number of propidium iodide puncta per EMO (n=10 EMOs per replicate). Statistical significance was measured for each replicate using a Student’s two-tailed t-test.

### Reconstitution of immune cell recruitment to *Chlamydia* infected EMOs

*Chlamydia* promotes the recruitment of immune cells, including neutrophils, which interact with and can invade infected epithelia to make direct contact with inclusions in infected animals (Rank et al., 2011). Their recruitment and activation at infection foci can damage the UGT (Lacy et al., 2011; Lee et al., 2010a; Lijek et al., 2018). However, *Chlamydia* has evolved strategies to limit the function of neutrophils. For example, the *Chlamydia* protease CPAF can inhibit neutrophil activation and netosis by cleaving neutrophil receptors (Rajeeve et al., 2018). Thus, we explored the application of the infection model by reconstituting aspects of the engagement of cellular immunity and its potential impact on *Chlamydia* infection. We co-cultured infected EMOs with primary bone marrow derived neutrophils and used high-resolution and time-lapse microscopy to quantify and visualize their recruitment to EMOs and interactions with infected epithelial cells.

EMOs were microinjected with mCherry-expressing *C. trachomatis* L2 or fluorescently labeled dextran. At 4 or 20 hours post-infection, primary fluorescently labelled neutrophils (CellTracker™) were added to the media for an additional 20 hours prior to imaging (Fig. 6A). Neutrophils infiltrated the Matrigel and migrated specifically to infected EMOs (Fig. 6A-B). EMOs microinjected with fluorescent dextran failed to recruit a significant number of neutrophils, indicating that the transient damage from the microinjection does not induce a chemoattracting response (Fig. 6B). We observed a subset of neutrophils make contact directly with infected EMO epithelial cells and even inclusions but did not appear to undergo netosis (Fig. 6C-D). Using timelapse spinning disk microscopy, we visualized neutrophil dynamics at infected EMOs and again observed a subset of neutrophils interact with the infected EMO and make contact with an intracellular inclusion (Fig. 6E-F).

**Figure 6.**
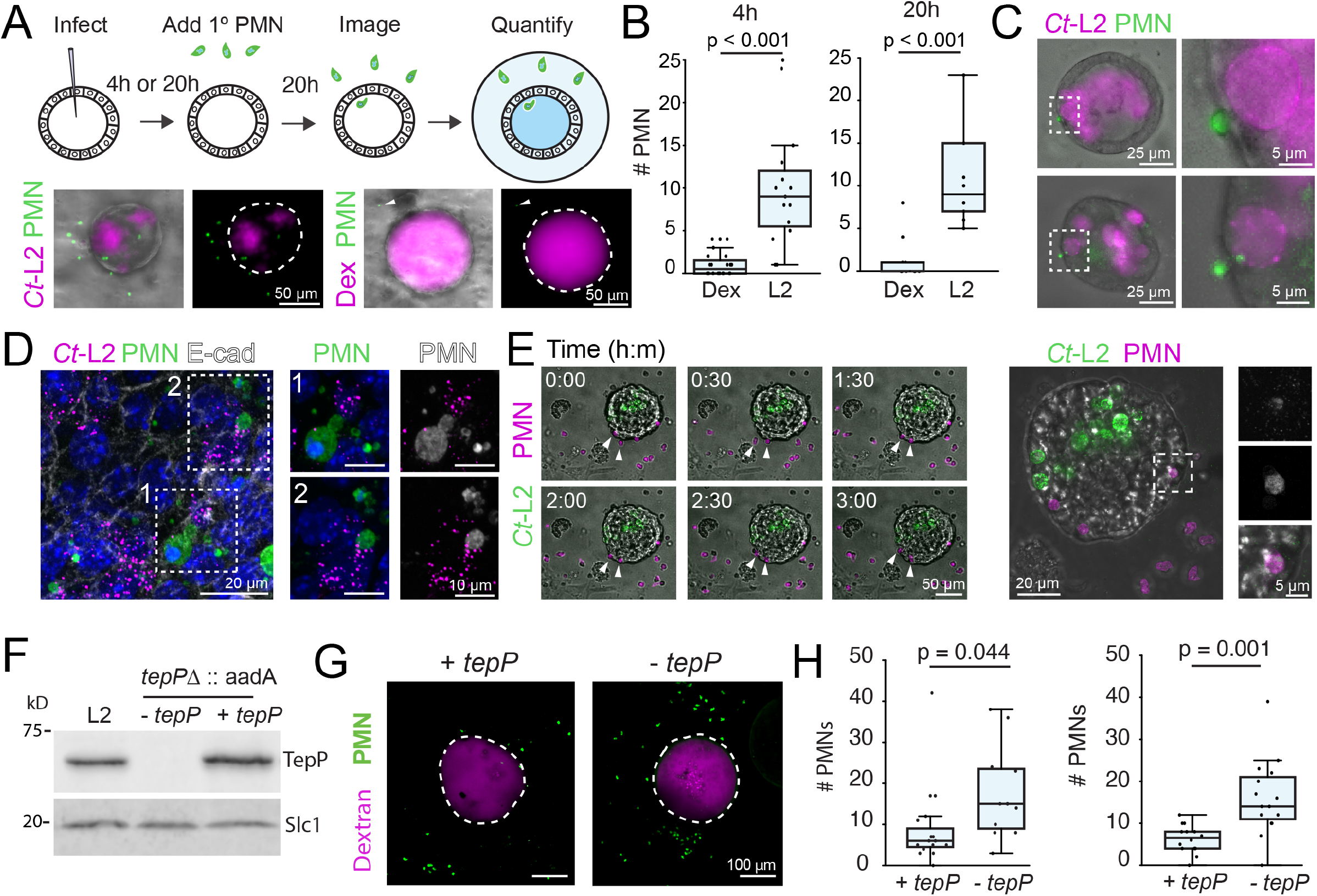
The recruitment of neutrophils to infected organoids is inhibited by the effector TepP. (**A**) Schematic representation of EMO infections and addition of neutrophils/polymorphonuclear (PMN) cells (top) and a representative widefield microscopy images (bottom) of EMOs infected with *Ct*-L2 or microinjected with Texas-Red dextran, cultured for 20 hours, and subsequently co-cultured with 5-chloromethylfluorescein diacetate (CFMDA)-labeled primary neutrophils for an additional 20 hours. (**B-D**) PMNs are specifically recruited to infected EMOs (B) and make contact with infected epithelial cells and intracellular inclusions (C-E). EMOs were infected with *Ct*-L2 for either 4 hours or 20 hours, co-cultured with PMNs for an additional 20 hours, and imaged by widefield microscopy. Quantification of PMN recruitment to infected EMOs at indicated times (4 hpi, n = 15-20 EMOs; or 20 hpi n = 10 EMOs) (B) and widefield microcopy images of tdTomato-expressing PMNs (green) contacting infected epithelial cells (C). High-resolution imaging of EMOs infected with mCherry-expressing *Ct*-L2 and cocultured with CFMDA-labeled PMNs. EMOs were stained for E-cadherin and imaged by confocal microscopy (D). Time-lapse microscopy of PMN recruitment to infected EMOs (**E**). EMOs were infected with GFP-expressing *Ct*-L2, co-cultured with tdTomato-expressing PMNs for 20 hours, and imaged by spinning disk confocal microscopy (E). Arrows denote PMNs moving towards and interacting with infected EMO. Maximum projection of higher magnification (*right panel*) of infected EMO. Enlarged box shows PMN co-localizing with *Ct*-L2 within the EMO. (**F-H**) TepP limits the recruitment of PMNs to infected EMOs. Whole cell lysates of wild type, *tepP* mutants or *tepP* mutants complemented with *tepP* on a plasmid were characterized by western blot analysis (F). The *Chlamydia* chaperone Slc1is shown as a loading control. EMOs were injected with a TepP mutant complemented with an empty vector (-*tepP*) or a vector expressing TepP (+ *tepP*) and Texas-Red dextran as marker of microinjection, co-cultured with CFMDA-labeled PMNs at 4 hpi, and imaged by 3D spinning disk confocal microscopy 20 hours later. Representative maximum projection images show PMNs recruited to infected EMOs (G) and the quantification of two independent experiments: left, n = at least 15 EMOs per infection; right, n=10 EMOs per infection) (H) are shown. Statistical significance was measured separately for each replicate in (Panels B, H) using a Student’s two-tailed t-test.

Infected epithelial cells secrete IL-6, IL-8, and members of the CXC chemokine family that could drive the recruitment of neutrophils towards *Chlamydia*-infected cells (Dessus-Babus et al., 2000; Frazer et al., 2011; Rasmussen et al., 1997). In mice, the absence of CXCR2, the CXC chemokine receptor, reduces acute inflammation and pathology in the UGT without affecting bacterial burden (Lee et al., 2010b). A transcriptional analysis of infected endocervical epithelial cells, indicated that the *Chlamydia* effector TepP enhances early type I interferon responses and dampens the expression of interleukin-6 (IL-6) and CXCL3 (Chen et al., 2014), chemokines that promote neutrophil chemotaxis (Fielding et al., 2008; Wright et al., 2014). We tested if TepP played a role in in regulating neutrophil infiltration by infecting EMOs by first generating a TepP knock out mutant by targeted insertion of an *aadA* cassette into the *tepP* locus (Fig. 6G). This mutant was then transformed with complementing plasmid expressing TepP under its native promote (+ *tepP*) or an empty vector (− *tepP*), and the resulting strains microinjected into EMOs followed by co-culturing with primary neutrophils at 4 hours post microinjection. Infected EMOS were imaged live for 20 hours. *C. trachomatis* strains lacking TepP showed significantly more neutrophil recruitment to the EMO, suggesting that TepP acts to dampen the innate immune response by limiting neutrophil influx (Fig. 6H-I).

Collectively, these data show that we can reconstitute the infiltration of innate immune cells to infected epithelia and address the role of *Chlamydia* virulence factors regulating the immune response.

## DISCUSSION

We developed a primary organoid model system to interrogate the cell biology *Chlamydia* infections in a cellular context that better mimics the architecture and diversity of cells present in the UGT epithelia, the target of these pathogens. Despite differences in the cytoskeletal and endomembrane organization in EMO cells compared to traditional two-dimensional tissue culture systems, we found remarkable conservation in the type of intracellular processes targeted by *Chlamydia* and in the effector proteins used. Our observations complement other recently reported models, including a fallopian tube organoid infection model (Kessler et al., 2019), and an endometrial organoid infection model that shows *Chlamydia* replicates within EMO epithelia and is amenable to inducible expression of *Chlamydia* Incs (Bishop et al. 2020, *In Press*). These models, however, do not infect the apical surfaces of intact organoids, which precludes the ability to monitor the natural route of *Chlamydia* invasion.

We provide evidence that *Chlamydia* reprograms the epithelial cytoskeleton and disrupts epithelial structure, at least transiently, during apical invasion and inclusion formation. These results are in agreement with infections in fallopian tube explants where *Chlamydia* infection disrupts epithelial architecture, altering the organization of polarity proteins in infected and uninfected neighboring cells (Kessler et al., 2012). Although we observe marked recruitment of β-catenin to early inclusions and *Chlamydia* entry sites, this localization does not occur in mid-to-late stage inclusions. Nevertheless, β-catenin promotes *Chlamydia* infection, possibly through Wnt-based signaling as pharmacological inhibition of Wnt signaling in transformed endometrial epithelial cells reduces *Chlamydia* growth (Kintner et al., 2017).

An advantage of this model lies in the ability to generate organoids from transgenic mouse lines. For instance, we used the ROSA^mTmG^ to better delineate single cell boundaries while following the dynamics of inclusions exiting the host cell. While all of the *C. muridarum* inclusions we observed exited apically, we observed a subset of *C. trachomatis* L2 inclusions positioned towards and exiting through the basolateral membrane, reminiscent of the observation in polarized endometrial cancer cells infected with *C. trachomatis* serovar E and treated with tamoxifen (Hall et al., 2011). It is unclear whether the exit strategy depends on the organization of polarized epithelial cell, cell-type specific responses, or serovar-specific functions. Because EMOs are comprised of a multiple epithelial subtypes (Fitzgerald et al., 2019), this model can be used to identify cell type-specific responses to infection.

We extended the application of EMOs to monitor the interaction between primary immune cells and infected epithelial cells. The rapid and specific recruitment of neutrophils to infected EMOs indicate that this model will be useful to dissect the dynamics of immune cells interaction with *Chlamydia* infected cells and the role played by host factors that regulate the innate immune responses. Neutrophils appear to have a limited role in clearing the infection, but rather influence the adaptive immune response by promoting the recruitment of T cells (Lacy et al., 2011). Studies using a combination of host and *Chlamydia* genetics can further identify the interaction between *Chlamydia* and the innate and adaptive immune response. For instance, we tested the effect of the T3S effector TepP on the influx of neutrophils to infected EMOs. Because neutrophil infiltration is regulated by type I and type III interferons (Blazek et al., 2015; Seo et al., 2011), TepP-dependent regulation of the interferon response and chemokine expression may function in part to reduce immune cell infiltrates and promote survival in the UGT.

In conclusion, we provide a high-resolution blueprint for the cell biology of *Chlamydia* infections in primary endometrial organoids. This foundation will provide a framework for future studies that target host and pathogens to better identify how *Chlamydia* subverts epithelial biology and the innate immune response.

## Materials and Methods

### Ethics statement

All animal experiments were approved and performed in accordance to the Duke University Institutional Animal Care and Use Committee.

### Cell lines and conditioned medium

Vero cells were purchased from ATCC (CCL-81) and cultured in Dulbecco’s Modified Eagle’s Medium (DMEM; Life Technologies) containing 10% fetal bovine serum (FBS; Sigma-Aldrich). L-WRN cells were purchased from ATCC (CRL-3276) and cultured in DMEM containing 0.5 mg/mL geneticin (Gibco) and 0.5 mg/mL hygromycin B (ThermoFisher) at 37°C with 5% CO2. Prior to generating conditioned medium, the cells were passaged twice in media without antibiotics. Conditioned medium was generated as previously described (Miyoshi and Stappenbeck, 2013). In brief, cells were plated in T175 tissue cultured treated flasks and grown to greater than 90% confluence. 25 mL of 1:1 DMEM/F12 (Gibco) media was collected every 24 hours, centrifuged at 500 x g for 5 min, and stored in 4°C. At the end of five days, the media were combined, sterile filtered, and stored in −80C.

### *Chlamydia* strains, propagation, and transformation

*C. trachomatis* strains L2/434/Bu (CTL2; ATCC VR-902B), the genetically-modified M007, M407, and M923 chemical mutants (Kokes et al., 2015; Sixt et al., 2017) were propagated in Vero cells, harvested at 44-48 hpi by water lysis, sonication, diluted in SPG (sucrose-phosphate-glutamate) buffer to 1x concentration (75 g/l sucrose, 0.5 g/l KH4PO4, 1.2 g/l Na2HPO4, 0.72 g/l glutamic acid, pH 7.5), and stored as single use aliquots at −80°C. The *C. muridarum* strain CM006 was a gift from Catherine O’Connell (University of North Carolina), and was propagated in Vero cells and harvested at 36 hpi as above. The CM006 strain was transformed with pNigg-SW2-GFP as follows: Approximately 10^7^ IFU were incubated with 10 μg DNA in buffer containing 0.9 mM calcium chloride for 30 min, added to confluent Vero cells in a six well plate, and centrifuged at 3,000 rpm for 30 min at 10°C. At 12 hours post-infection, 1 U/mL penicillin was added. The infections were passaged every 36 hours until inclusions were present and fluorescent. Transformants were subsequently plaque-purified to obtain a clonal strain.

To generate the spectinomycin-resistant TepP-deficient *C. trachomatis*, CTL2 was transformed using the TargeTron gene disruption system (Sigma-Aldrich) as the previously described TepP mutant (Carpenter et al., 2017) with the exception that an *aadA* cassette was inserted into the targeting site (between amino acids 821-822). Transformants were expanded in Vero cells in the presence of 150 ug/mL spectinomycin as described above, plaque purified, and verified by PCR analysis using primers flanking the insertion site and the *aadA* cassette. The *ΔtepP::aadA* strain was complemented with the *E.coli-Chlamydia* shuttle vector p2TK2-SW2 (Agaisse and Derré, 2013) containing a *bla* cassette and the TepP open reading frame containing its upstream promoter sequence. In brief, 10^8^ IFU were incubated with 10 μg DNA in buffer containing 0.9 mM calcium chloride for 30 min, added to confluent Vero cells in a six well plate, and centrifuged at 3,000 rpm for 30 min at 10°C. At 12 hours post-infection, 1 U/mL penicillin was added. The infections were passaged every 48 hours until inclusions were present where penicillin concentration was increased to 10 U/mL. Transformants were subsequently plaque-purified in the presence of both spectinomycin and penicillin to obtain a clonal strain and verified by PCR using primers targeting both the *aadA* and *bla* cassette and the full-length *tepP* sequence.

### Western Blots

For western blots, Vero cells were infected with the indicated strains for 48 hours, water lysed, sonicated, and boiled in Laemelli buffer for 10 min. The lysates were centrifuged at 9,600 x g for 5 min and equal volumes were resolved using a 10% SDS-PAGE gel. The resolved proteins were transferred to a nitrocellulose membrane (Amersham), blocked with a PBS solution containing 5% milk for one hour at 25°C, and sequentially probed with primary anti-TepP (1:500; (Chen et al., 2014)) and anti-Slc1 (1:1000; (Chen et al., 2014)) and the LiCOR infrared-conjugated secondary antibodies (1:10,000). Membranes were imaged using an Odyssey LiCOR.

### EMO generation from the mouse endometrium and hormone stimulation

The C57/BL6J and B6.129(Cg)-Gt(ROSA)26Sor^tm4(ACTB-tdTomato,-EGFP)Luo/J (Strain no. 007676) mouse strains were purchased from Jackson Laboratories. The ZO-1 GFP knock-in mouse line was a generous gift from Dr. Terry Lechler (Duke University). The endometrium from ~ 6-8 week old females were dissected, washed in cold Dulbecco’s phosphate-buffered saline (PBS; Gibco) on ice, cleaned of vascular and fat tissue, minced into ~ 2 mm pieces, transferred to DMEM (Gibco) containing 0.2% collagenase A (Thermo), 10% FBS, and 1U/mL penicillin/streptomycin, (Gibco) then incubated on a shaker at 37°C for 2.5 – 3 hours. The tissue was washed three times with cold PBS and mechanically disrupted by shaking in cold 10 mL PBS containing 0.1% bovine serum albumin (BSA; Fraction V, Equitech-Bio) for 1 min. After allowing the tissue to settle for one minute, the supernatant was collected and passed through a 70 μm strainer (Falcon). The mechanical disruption was repeated once more, and the strainer was subsequently inverted over a new 50 mL conical tube and the epithelia were washed off the strainer three times with 10 mL PBS containing 0.1% BSA. The epithelial fraction was centrifuged at 500 x g for 5 min at 10°C, resuspended in cold DMEM/F12 (Gibco), mixed with low-hormone Matrigel (Corning) at a 1:1 ratio, and pipetted in 35 μL drops in a 24 well plate or 125 μL drops in a 35 mm glass-bottom dish (for microinjections). The plates were incubated at 37°C for 40 min before gently overlaying 2 mL 1:1 DMEM/F12 media containing 50% L-WRN conditioned media, 50 μg/mL gentamicin (Gibco), and 50 ng/mL EGF (StemCell Technologies). The media was changed every 2-3 days. For hormonal stimulation studies, EMOs were cultured in the presence of 17-β-estradiol (E2; Sigma-Aldrich) for four days.

### Organoid microinjection

Microinjections were performed using an Eppendorf FemtoJet 4x coupled with a Stereo Microscope (Nikon). *Chlamydia* strains were diluted in PBS to a final concentration of 5E^5^-5E^6^ IFU, vortexed for 30s, and pipetted into a glass needle. Organoids were punctured once using a steep vertical angle and manually injected with equal volumes. When organoids were injected alone or co-injected with 3kD Texas-Red dextran (Invitrogen), dextran was used at a final concentration of 0.01 mg/mL.

### Primary neutrophil isolation, co-culture with infected EMOs

To isolate primary neutrophils, mouse femurs were dissected, cleaned with an ethanol-soaked Kim wipe to remove other tissue, dipped in 70% ethanol then twice in RPMI media (Gibco). Both ends of the femur were cut and 10 mL RPMI media was passed through the bone with a 25G needle (BD Biosciences). The supernatant was centrifuge at 600 x g for 5 minutes, resuspended in 5 mL red cell lysis buffer (Millipore), and incubated for 5 minutes. An additional 5 mL of RPMI media was added to the cells and centrifuged at 600 x g for 5 minutes. The cells were resuspended gently in 0.5 mL RPMI media before following the negative selection EasySep™ Mouse Neutrophil Enrichment Kit (StemCell Technologies). Neutrophils were labeled with 5 μM CellTracker (CFMDA; ThermoFisher) for 10 minutes at 37°C, centrifuged at 300 x g for 10 minutes at 4°C, and resuspended in 0.5 mL cold RPMI. Neutrophils were counted and 3E^5^ cells were added directly to the media and cultured for an additional 20 hours.

### Antibodies

For immunofluorescence microscopy, the following antibodies were used: mouse anti-GM130 (BD Biosciences, Cat no. 610822; 1:200), anti-alpha-tubulin (Sigma, Cat no. T5618, Clone B-5-1-2; 1:200), anti-Muc1 (Cell Signaling Technologies, Cat no. VU4H5; 1:100), anti-β-catenin (BD Biosciences, Cat no. 610153 Clone 14; 1:400), anti-MOMP (Santa Cruz, Cat no. 57678; 1:500), anti-pan-Keratin (Sigma-Aldrich Cat no. C2562; 1:100), and rabbit anti-E-cadherin (Cell Signaling Technologies; Cat no. 3195S, Clone 24E10; 1:200), anti-acetylated-alpha-tubulin (Cell Signaling Technologies, Cat no. 5335S 1:200), anti-SEPT2 (Protein Tech; Cat no. 11397-1-AP; 1:100), anti-Cap1 (Gift from A. Subtil; 1:250).

### Brightfield and immunofluorescence microscopy

Organoid growth was monitored by brightfield microscopy using the EVOS FL Cell Imaging System (ThermoFisher) equipped with 2x/0.06 and 10x/0.25 NA objectives and a CCD camera. For fixed samples, organoids were rinsed twice with warm PBS and incubated with warm PBS containing 3% formaldehyde (Sigma) for 20 minutes. The fixative was removed gently, and the organoids were resuspended in 0.25% ammonium chloride (Sigma-Aldrich) and transferred to a 1.5 mL tube, centrifuged at 500 x g for 5 min, resuspended gently in 2% BSA (Sigma-Aldrich) containing 0.1% triton x-100 and incubated with gentle rocking for 30 minutes. Organoids were centrifuged again at 500 x g for 5 min, incubated with the indicated primary antibodies diluted in 0.5 mL 2% BSA containing 0.1% triton x-100, and incubated at 25°C for 2-3 hours or overnight at 4°C with gentle rocking. Organoids were washed once with 2% BSA, centrifuged at 500 x g for 5 min, and incubated with secondary antibodies diluted in 1.0 mL 2% BSA containing 0.1% triton x-100 for 1.5 hours at 25°C with gentle rocking. Acti-stain conjugated to Alexa-555 (1:500; Cytoskeleton, Inc) and Hoechst (2 μg/mL; ThermoScientific) were added for the final 20 minutes of incubation with the secondary antibodies. The organoids were centrifuged at 500 x g for 5 min and resuspended in 30 μL Vectashield (Vector Labs; H-1000) using a cut pipet tip and pipetted onto a coverslide. A coverslip was overlaid gently and sealed with nail polish.

Organoids were imaged using an inverted confocal laser scanning microscope (LSM 880; Zeiss) equipped with a motorized stage, Airyscan detector (Hamamatsu), and diode (405 nm), argon ion (488 nm), double solid-state (561 nm), and helium-neon (633 nm) lasers. Images were acquired using a 20x/0.8 NA air or 40x/1.2 NA water objective (Zeiss) and deconvolved using automatic Airyscan Processing in the Zen Software (Zeiss). Images were opened in ImageJ (NIH) or the Imaris software and exported TIFFs were rendered in the Adobe suites (Photoshop and Illustrator). Only linear adjustments were made to fluorescence intensity.

### Time-lapse microscopy of inclusion dynamics and neutrophil recruitment

Organoids cultured in 35 mm glass bottom dishes (Mat-Tek) were imaged live using an inverted microscope (Zeiss AxioObserverZ.1) equipped with a motorized stage containing a heated Insert P environmental chamber (Zeiss), XLIGHT V2 spinning disk unit (Crest Optics), and an ORCA Flash 4.0 V3 camera (Hamamatsu). Images were acquired using a 20x/0.8 NA objective (Olympus), an LDI multiline laser (89 North) using two micron sections at 6-10 minute intervals for 16-18 hours. Using the same microscope, neutrophil recruitment to infected EMOs was imaged using five micron confocal sections and spanning ~ 100 μm above and below each organoid. Timelapse microscopy of neutrophil motility was performed using a 20x/0.8 NA objective (Olympus) or a 60x/1.4 NA objective (Olympus). Confocal sections (2 μm) were acquired every three minutes for the duration of three hours. All images were rendered in ImageJ (NIH). Exported TIFFs were reconstructed in the Adobe suites (Photoshop, Illustrator).

### Image analysis

To quantify EMO size, microscopy images were imported into ImageJ and converted to 8-bit TIFFs. EMO borders were identified using the find edges algorithm and converted to a binary image. Using the find maxima algorithm, each EMO was identified, the area exported into Excel and plotted in R.

Golgi reorganization around the inclusion was quantified in ImageJ. Microscopy images were imported and converted to 8-bit TIFFs. The inclusion edges were manually traced to measure the perimeter length. Using the segmented line tool, the Golgi signal at the inclusion was also traced to measure its length. Each value was imported into Excel to generate the percent distribution around the inclusion perimeter and subsequently plotted in R.

To quantify cell death, microscopy images of EMOs incubated with propidium iodide were quantified in ImageJ. Microscopy images were imported and converted to 8-bit TIFFs. Maximum projection images were background subtracted using the rolling ball radius (value = 50) and puncta were identified using the find maxima algorithm.

Neutrophil recruitment to infected EMOs was quantified in ImageJ. Maximum projection images were background subtracted as above. To identify proximal neutrophils, a circle was placed around the inclusion 200 μm from the EMO basolateral edge. The number of neutrophils were identified using the find maxima algorithm, imported into Excel and plotted in R.

## Acknowledgements

We thank Dr. Lisa Cameron at the Duke Light Microscopy Core Facility and Dr. David Tobin for assistance with microscopy, and members of the Valdivia lab for critical feedback on this project. Dr. Jorn Coers and Dr. Ryan Finethy for assistance with mice and immune cell isolations, Dr. Terry Lechler for the ZO-1 GFP knock-in mice, and Dr. Agathe Subtil for the Cap1 antibody.

---

**Supplemental Figure 1.**
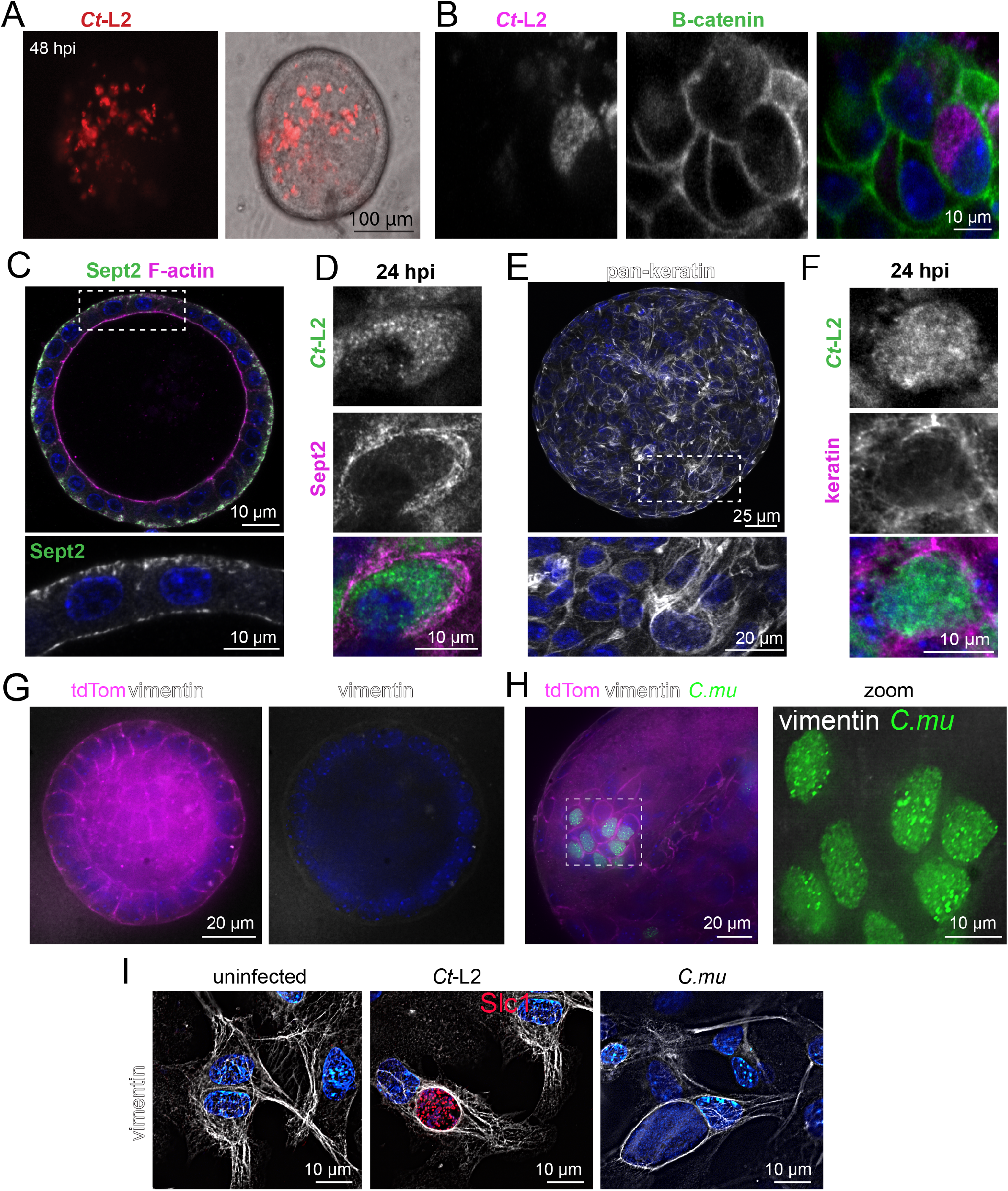
Mid-cycle localization of cytoskeletal elements to the *C. trachomatis* inclusion. **(A)** *Chlamydia* inclusions at 48 hours post-infection. Widefield microscopy images of EMO infected with mCherry-expressing Ct*-*L2. (B) β-catenin localization with respect to the *C. trachomatis* inclusion. Confocal microscopy images of an EMO infected with mCherry-expressing *Ct*-L2 for 48 hours and stained for β-catenin. **(C-D)** Septins localize to the *C. trachomatis* inclusion. Confocal microscopy images of an uninfected (C) and infected (D) EMO stained for Sept2 and F-actin **(E-F)** Keratins localize to the *C. trachomatis* inclusion. Confocal microscopy of an uninfected (E) and infected (F) EMO stained for cytokeratins and F-actin. **(G-H)** Vimentin is not expressed in EMO epithelia. (G) Widefield deconvolution image of an endometrial organoid expressing tdTomato and stained for vimentin. (H) EMO infected with *C. muridarum* for 24 hours and stained for vimentin. **(I)** Vimentin localizes to the *Chlamydia* inclusion in primary stromal fibroblasts. Widefield deconvolution images of primary stromal fibroblasts uninfected or infected with *C. trachomatis* or *C. muridarum* for 24 hours and stained for vimentin. For all images DNA stains used 2 μg/mL Hoechst (blue).

